## Supplementary material for "Countering reproducibility issues in mathematical models with software engineering techniques: A case study using a one-dimensional mathematical model of the atrioventricular node": Data Supplement

### 1 Supplementary Figures

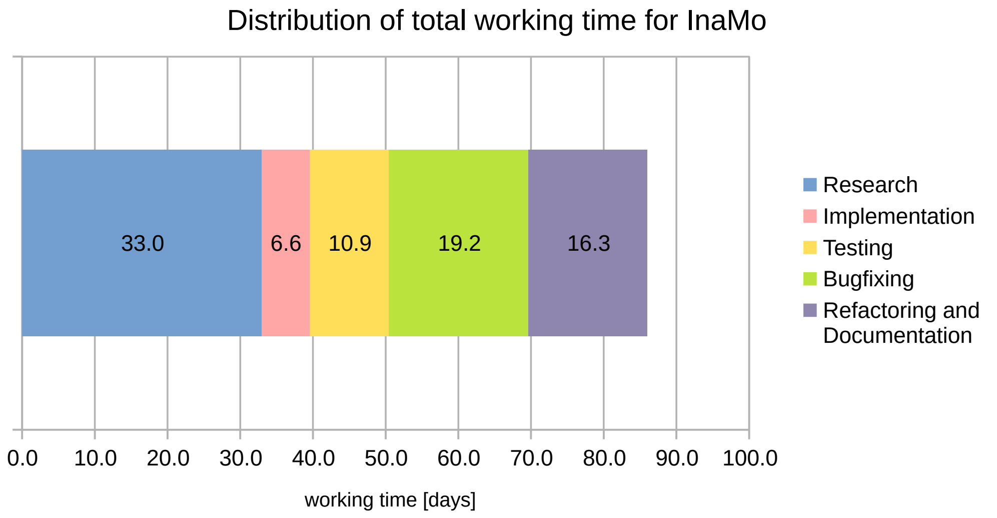

**Figure 1.** Estimation of working time distribution for the development of the Modelica version of the Inada model. Estimates were taken from time stamps of PDF comments in the articles listed in Figure 1 and of commits in the version control system Git. In a first step, we always assumed that a full day was spent on literature research or on the code in the repository. To correct for overlap on days where we worked both on the repository and added comments in the PDF documents, we scaled the whole dataset to the amount of unique days spend for development (86).

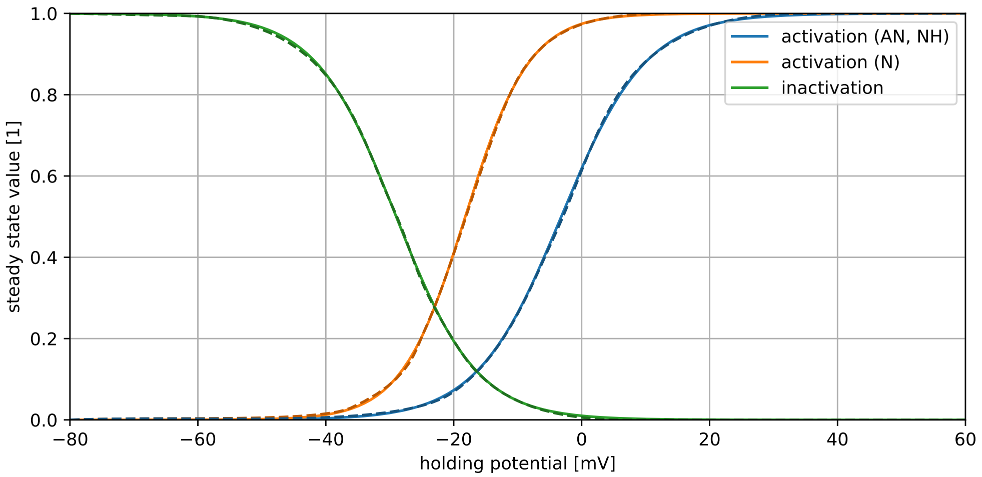

**Figure 2.** Steady states of gating variables in $I_{Ca,L}$. Solid lines: Simulation result of LTypeCalciumSteady. Dashed lines: Reference data extracted from Figure S1A and S1B in [1]. The plots are in perfect agreement. This figure was created with InaMo version 1.4.2, which is available under the DOI 10.5281/zenodo.4533008.

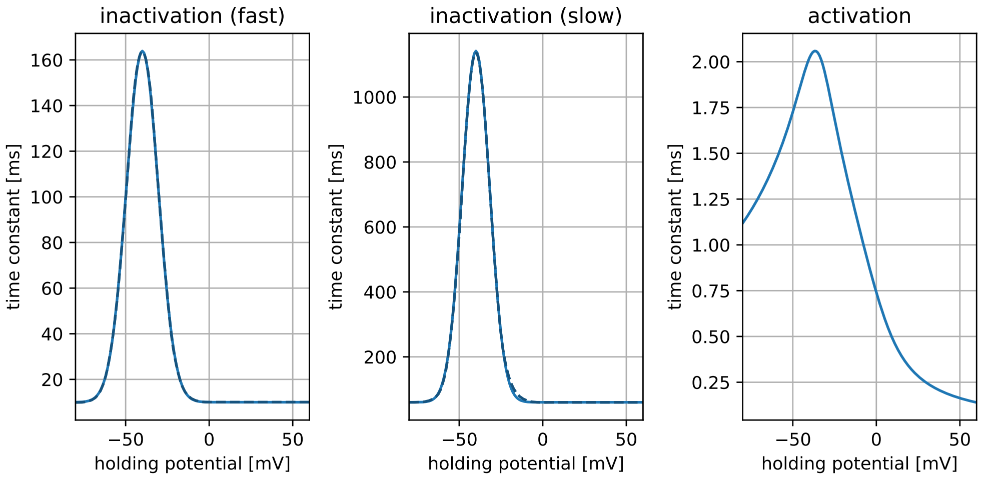

**Figure 3.** Time constants of gating variables in $I_{Ca,L}$. Solid lines: Simulation result of LTypeCalciumSteady. Dashed lines: Reference data extracted from Figure S1C (fast inactivation) and S1D (slow inactivation) in [1]. A reference plot for the activation gate is not provided in [1]. The plots are in perfect agreement. This figure was created with InaMo version 1.4.2, which is available under the DOI 10.5281/zenodo.4533008.

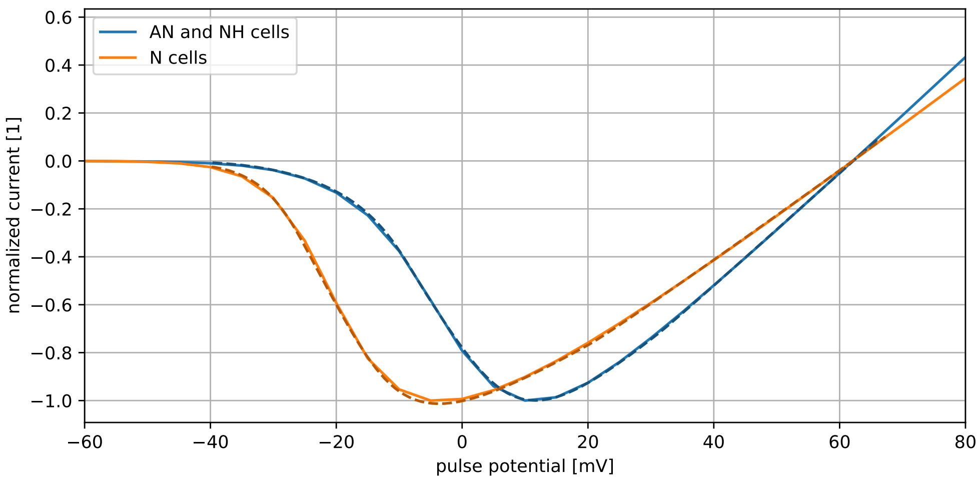

**Figure 4.** Reference plots for current-voltage relationship of $I_{Ca,L}$ obtained with a voltage pulse protocol with a holding potential of -70 mV, a holding duration of 5 s, and a pulse duration of 300 ms. Solid lines: Simulation result of LTypeCalciumIV. Dashed lines: Reference data extracted from Figure S1E in [1]. The plots are in perfect agreement. This figure was created with InaMo version 1.4.2, which is available under the DOI 10.5281/zenodo.4533008.

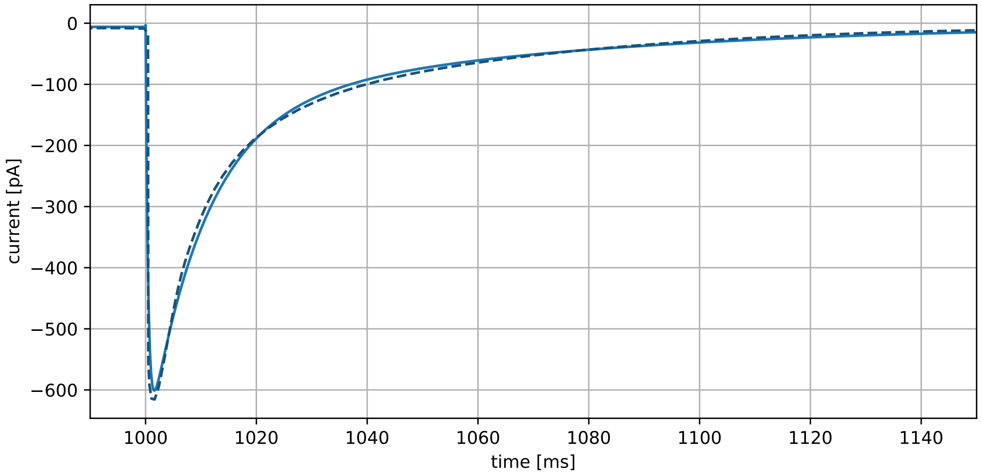

**Figure 5.** Time course of $I_{Ca,L}$ when switching from holding potential of -40 mV to prolonged stimulation at +10 mV. Solid line: Simulation result of LTypeCalciumStep. Dashed line: Reference data extracted from Figure S1H in [1]. The plots are in perfect agreement. Note, however, that while Inada *et al.* state that they used AN cells for plot S1H, NH cells had to be used instead to reach this agreement. Additionally, it seems that the x axis of Figure S1H is scaled differently than the measuring strip in the plot suggests. To obtain a good fit, time stamps extracted from the figure have to be multiplied by a scaling factor of 0.75. This figure was created with InaMo version 1.4.2, which is available under the DOI 10.5281/zenodo.4533008.

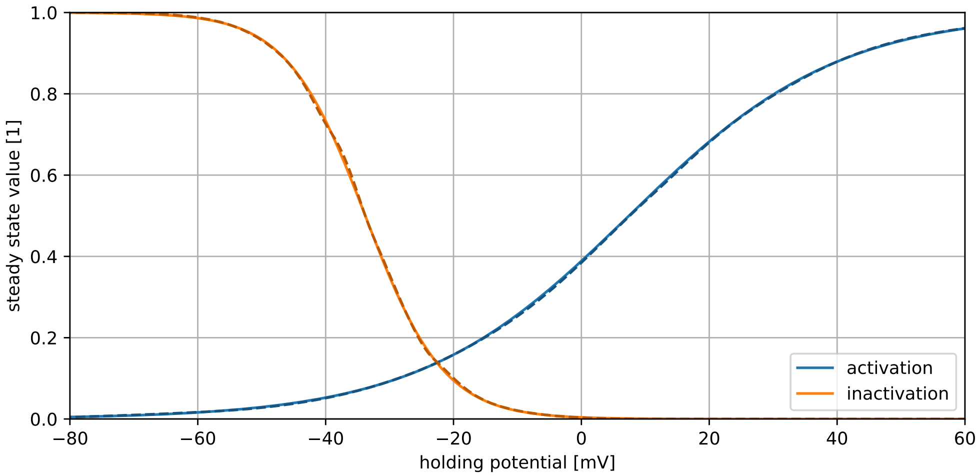

**Figure 6.** Steady states for activation and inactivation gates of $I_{to}$. Solid lines: Simulation result of TransientOutwardSteady. Dashed lines: Reference data extracted from Figures S2A and S2B in [1]. The plots are in perfect agreement. This figure was created with InaMo version 1.4.2, which is available under the DOI 10.5281/zenodo.4533008.

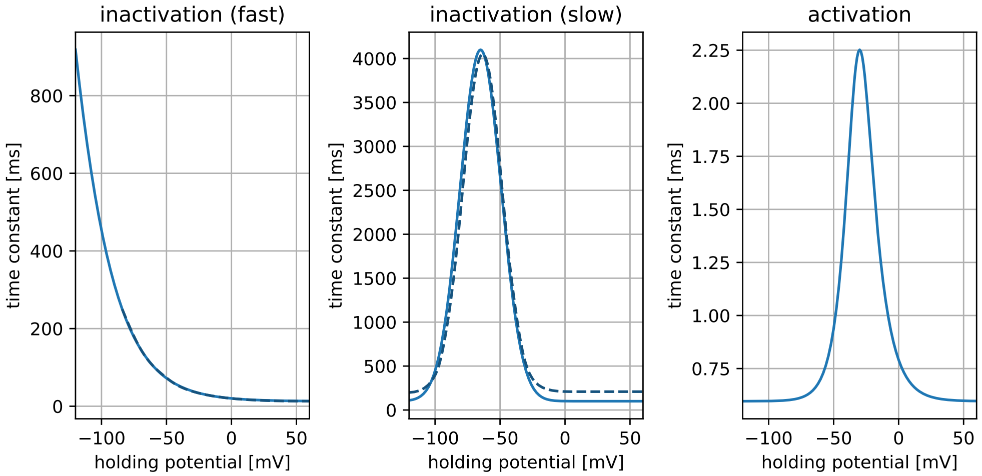

**Figure 7.** Time constants for gating variables in $I_{to}$. Solid lines: Simulation result of TransientOutwardSteady. Dashed lines: Reference data extracted from Figures S2C and S2D in [1]. A reference plot for activation is not provided in [1]. The fast inactivation is in perfect agreement, but the slow inactivation shows a lower minimum value. It seems that the minimum in Figure S2D is closer to 0.2 s than 0.1 s as given in [1] and the C++ code. Since article and C++ code agree on the value 0.1 s, we assume that Figure S2D was generated with an older version of the model. This figure was created with InaMo version 1.4.2, which is available under the DOI 10.5281/zenodo.4533008.

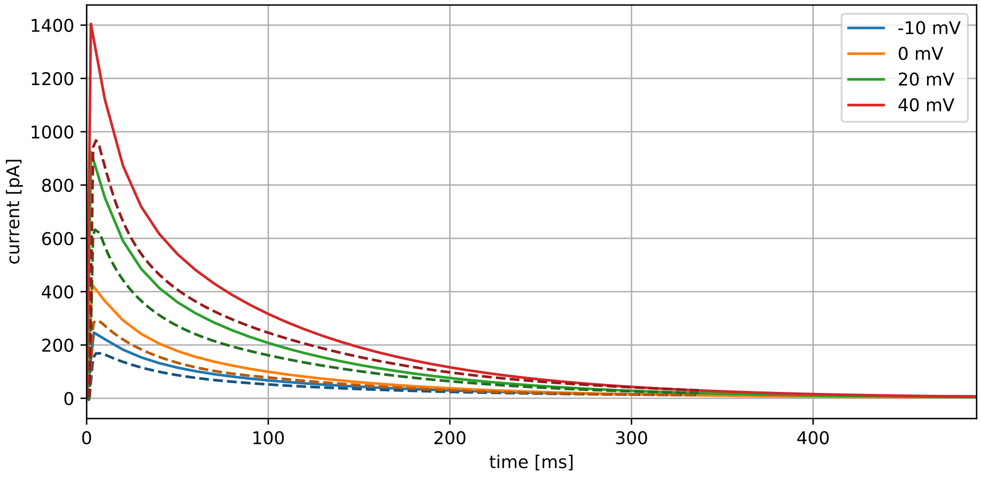

**Figure 8.** Time course of $I_{to}$ after a stimulation to different voltages for 500 ms from a holding potential of -80 mV that was kept for 20 s before the stimulation. Solid lines: Simulation result of TransientOutwardIV. Dashed lines: Reference data extracted from Figure S2E in [1]. While Inada *et al.* state that they used AN cells for plot S2E, NH cells were used instead to reach a better agreement to the reference plot. The absolute values for the current are larger for InaMo than for the reference. This may be due to differences in the holding duration, which was not reported by Inada *et al.*. We chose a holding duration 20 s for a full return to the steady state, since otherwise the previous pulse would influence the following. The differences to the reference vanish when a scaling factor of 0.75 is applied to the simulation results. This figure was created with InaMo version 1.4.2, which is available under the DOI 10.5281/zenodo.4533008.

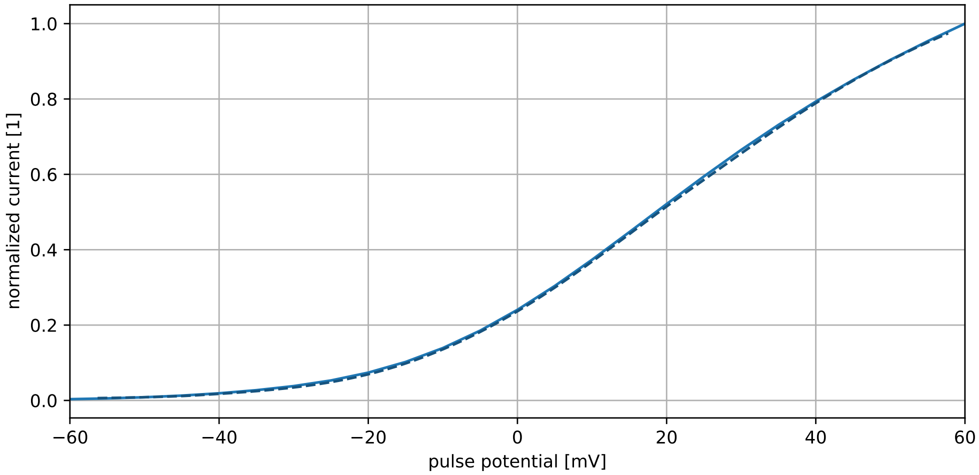

**Figure 9.** Current-voltage relationship of $I_{to}$ obtained with a voltage pulse protocol with a holding potential of -80 mV, a holding duration of 20 s, and a pulse duration of 500 ms. Solid line: Simulation result of TransientOutwardIV. Dashed line: Reference data extracted from Figure S2F in [1]. The plots are in perfect agreement. This figure was created with InaMo version 1.4.2, which is available under the DOI 10.5281/zenodo.4533008.

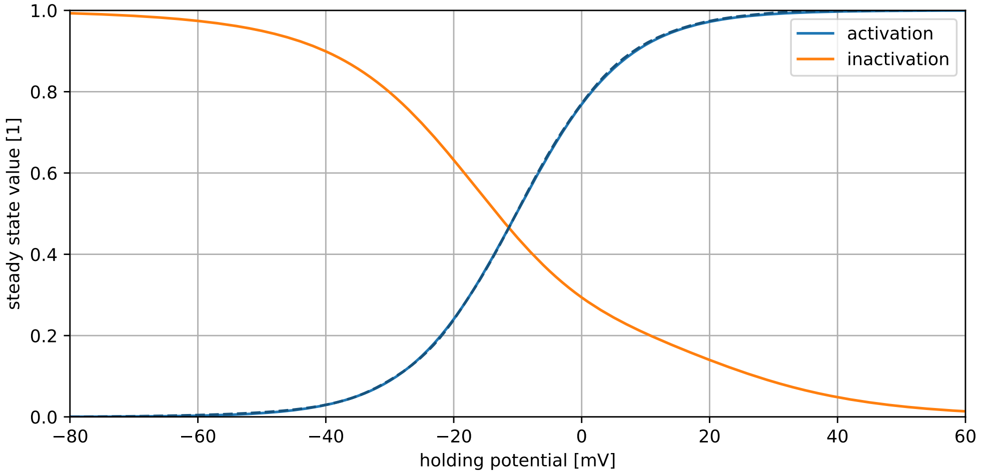

**Figure 10.** Steady state of the gating variables for $I_{K,r}$. Solid lines: Simulation result of RapidDelayedRectifierSteady. Dashed lines: Reference data extracted from Figure S3A in [1]. The plots are in perfect agreement. This figure was created with InaMo version 1.4.2, which is available under the DOI 10.5281/zenodo.4533008.

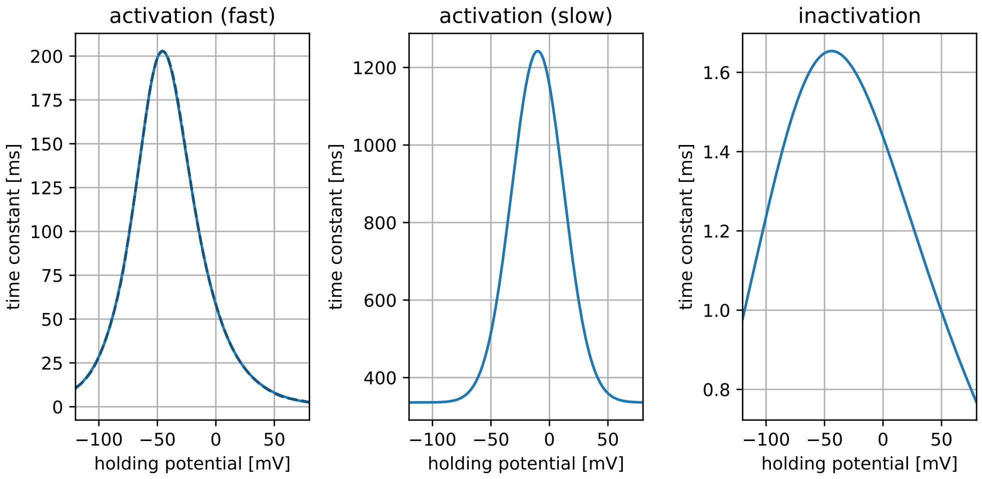

**Figure 11.** Time constants of the gating variables for $I_{K,r}$. Solid lines: Simulation result of RapidDelayedRectifierSteady. Dashed lines: Reference data extracted from Figure S3B in [1] (showing fast activation). Reference plots for activation are not provided in [1]. The plots are in perfect agreement. This figure was created with InaMo version 1.4.2, which is available under the DOI 10.5281/zenodo.4533008.

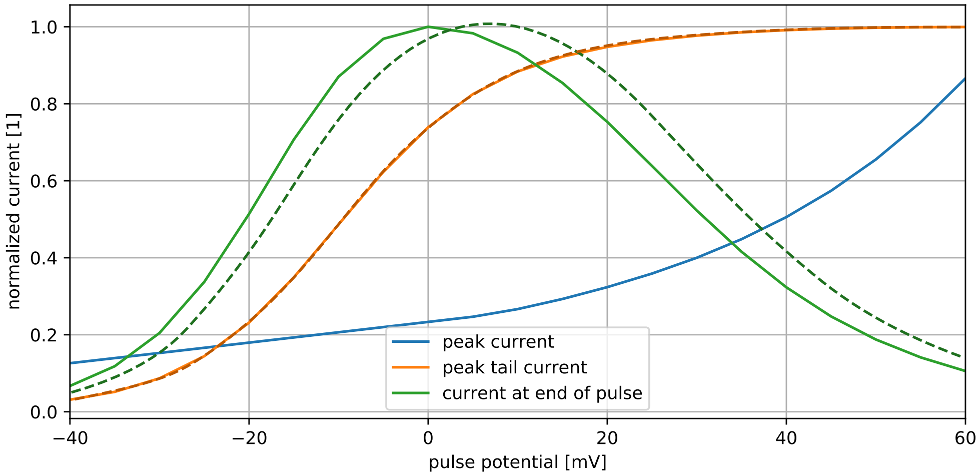

**Figure 12.** Current-voltage relationship of $I_{K,r}$ obtained with a voltage pulse protocol with a holding potential of -40 mV, a holding duration of 5 s, and a pulse duration of 500 ms. Solid lines: Simulation result of RapidDelayedRectifierIV. Dashed lines: Reference data extracted from Figures S3C and S3D in [1]. Figure S3C shows the current at the end of each stimulation pulse. Figure S3D shows the peak current obtained after the voltage shifts back from the pulse voltage to the holding potential. Data from Figure S3D is in perfect agreement with simulation results, but data from Figure S3C is shifted by 5 mV towards higher voltages. This could be explained if Inada *et al.* accidentally associated currents to the newly started pulse right after the current was measured instead of the previous pulse. This figure was created with InaMo version 1.4.2, which is available under the DOI 10.5281/zenodo.4533008.

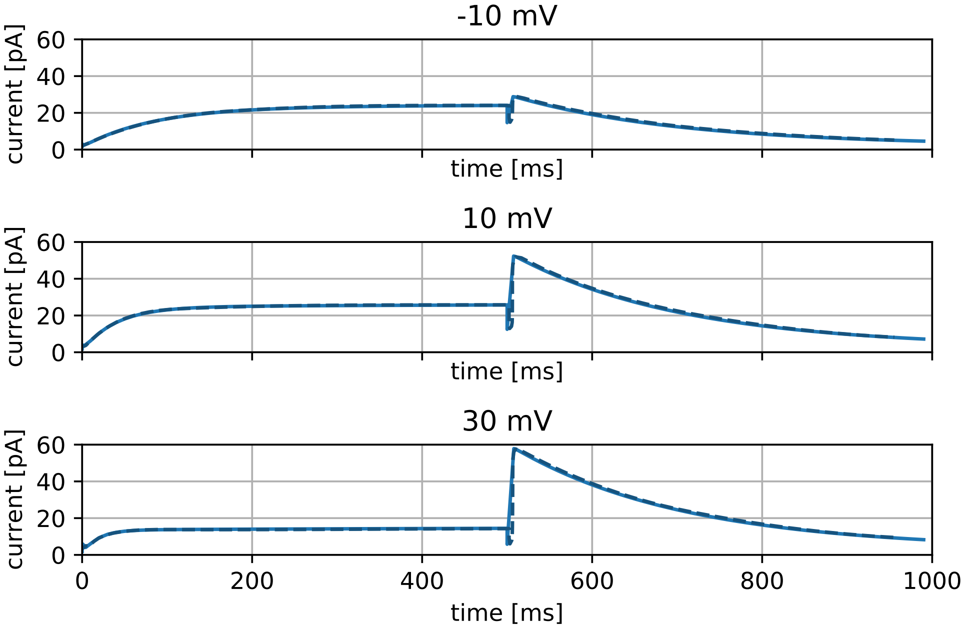

**Figure 13.** Current time course of $I_{K,r}$ after 500 ms pulses with different voltages from a holding potential of -40 mV which is held for 5 s. Solid lines: Simulation result of RapidDelayedRectifierIV. Dashed lines: Reference data extracted from Figure S3E in [1]. The plots are in perfect agreement. This figure was created with InaMo version 1.4.2, which is available under the DOI 10.5281/zenodo.4533008.

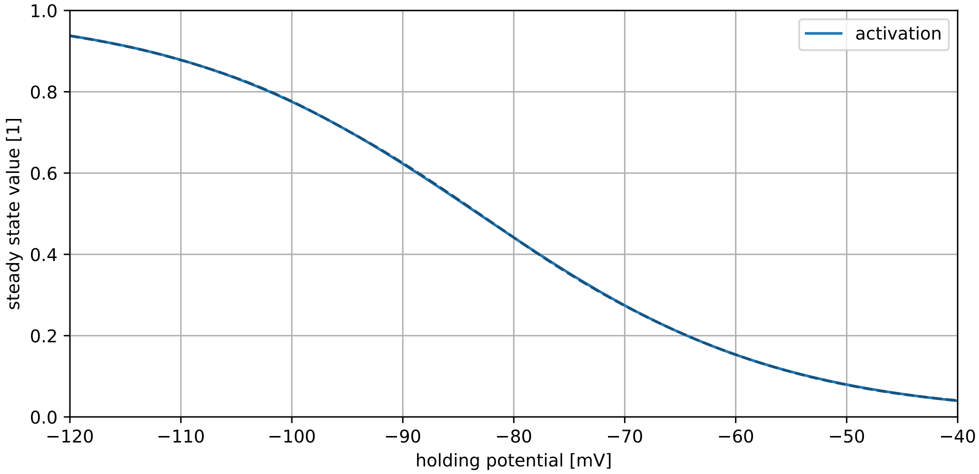

**Figure 14.** Steady state of activation gating variable in $I_f$. Solid line: Simulation result of HyperpolarizationActivatedSteady. Dashed line: Reference data extracted from Figure S4A in [1]. The plots are in perfect agreement. This figure was created with InaMo version 1.4.2, which is available under the DOI 10.5281/zenodo.4533008.

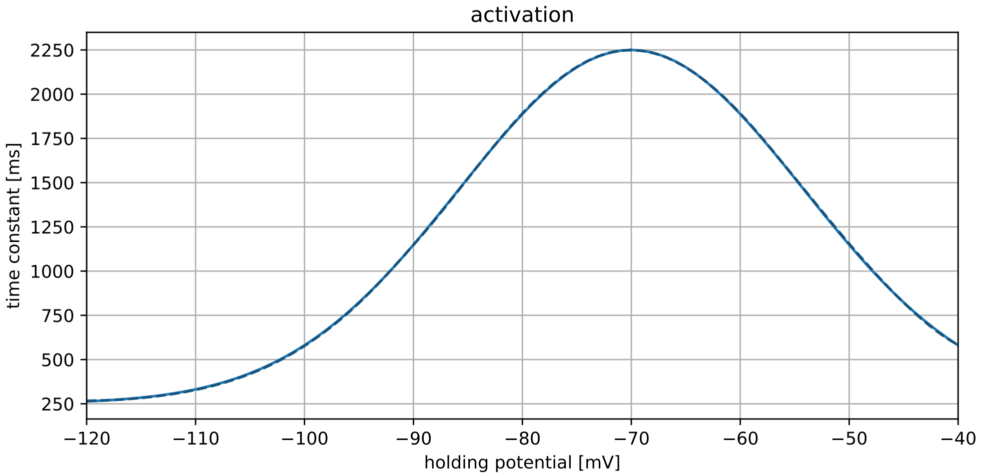

**Figure 15.** Time constant for activation gating variable in $I_f$. Solid line: Simulation result of HyperpolarizationActivatedSteady. Dashed line: Reference data extracted from Figure S4B in [1]. The plots are in perfect agreement. This figure was created with InaMo version 1.4.2, which is available under the DOI 10.5281/zenodo.4533008.

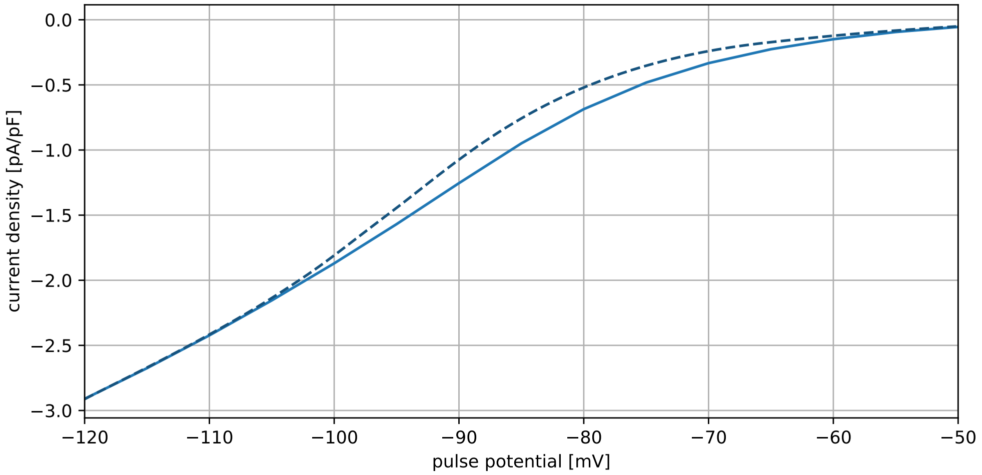

**Figure 16.** Current-voltage relationship of $I_f$ obtained with a voltage pulse protocol with a holding potential of -50 mV, a holding duration of 20 s, and a pulse duration of 4 s. Solid line: Simulation result of HyperpolarizationActivatedIV. Dashed line: Reference data extracted from Figure S4C in [1]. The plots diverge for pulse voltages between -100 to -60 mV with higher current densities in the reference data. We have no good explanation for this difference, especially considering that Figure 17, which covers similar voltage ranges using the same data, shows a perfect agreement for $I_f$. This figure was created with InaMo version 1.4.2, which is available under the DOI 10.5281/zenodo.4533008.

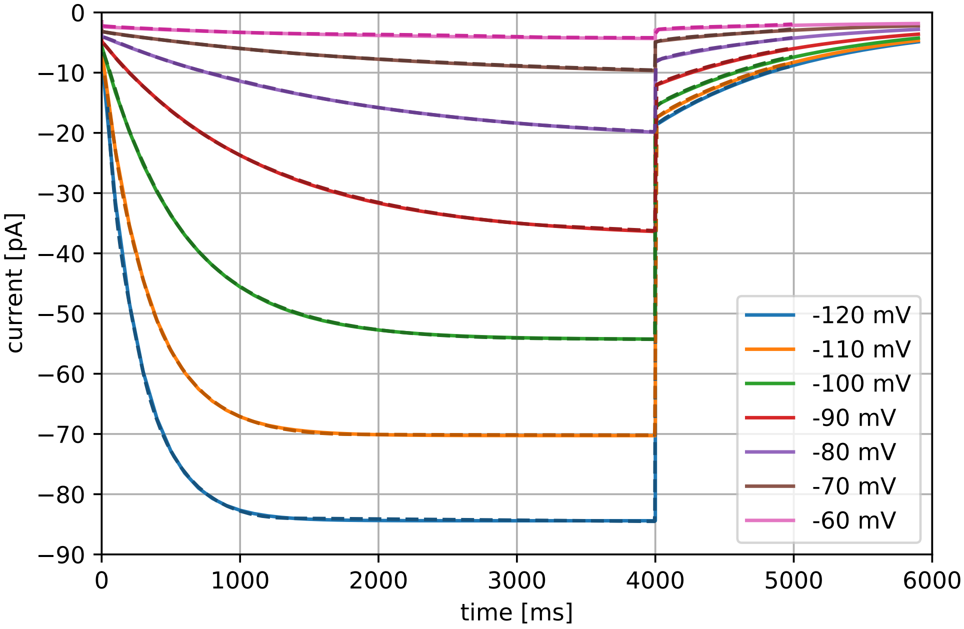

**Figure 17.** Time course of $I_f$ after 4 s of stimulation to different voltages from a resting potential of -50 mV, which was held for 20 s. Solid lines: Simulation result of HyperpolarizationActivatedIV. Dashed lines: Reference data extracted from Figure S4D in [1]. The plots are in perfect agreement. This figure was created with InaMo version 1.4.2, which is available under the DOI 10.5281/zenodo.4533008.

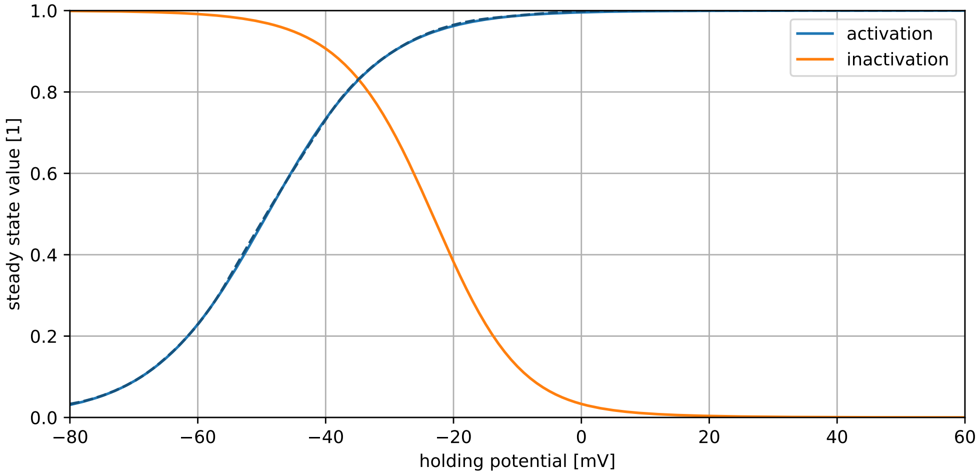

**Figure 18.** Steady state of gating variables in $I_{st}$. Solid lines: Simulation result of SustainedInwardSteady. Dashed lines: Reference data extracted from Figure S5A in [1]. A reference plot for inactivation is not provided in [1]. The plots are in perfect agreement. This figure was created with InaMo version 1.4.2, which is available under the DOI 10.5281/zenodo.4533008.

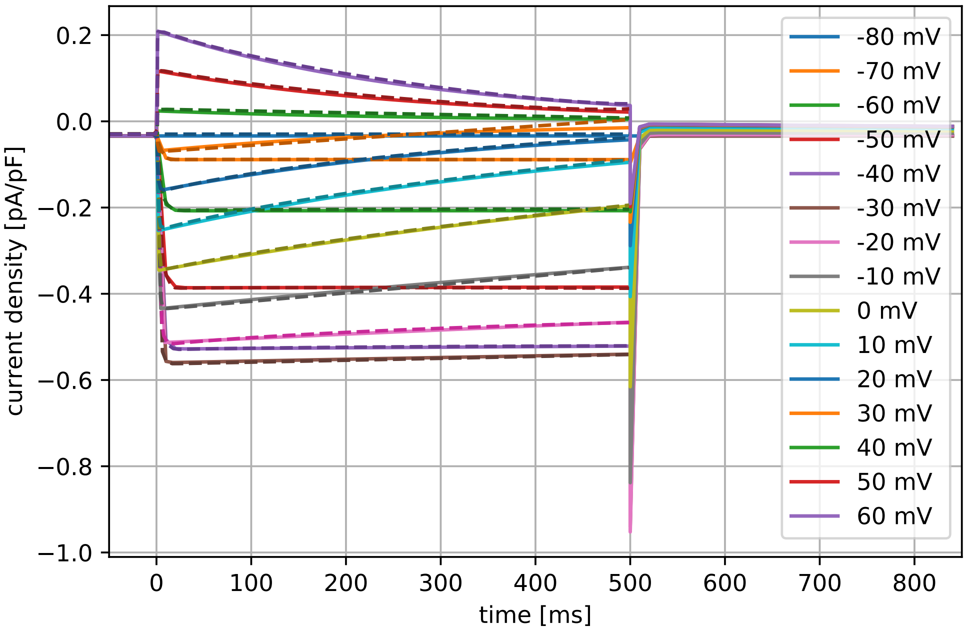

**Figure 19.** Current density time course of $I_{st}$ during a voltage pulse protocol with a holding potential of -80 mV, a holding duration of 15 s, and a pulse duration of 500 ms. Solid lines: Simulation result of SustainedInwardIV. Dashed lines: Reference data extracted from Figure S5B in [1]. We had to change the parameter $g_{st}$ to 0.27 nS instead of the default 0.1 nS reported in [1] in order to obtain a good fit to the reference data. Evidence that this is not due to an error in our implementation is provided by the good agreement between our results and those of [2] in Figures 29 and 30. This figure was created with InaMo version 1.4.2, which is available under the DOI 10.5281/zenodo.4533008.

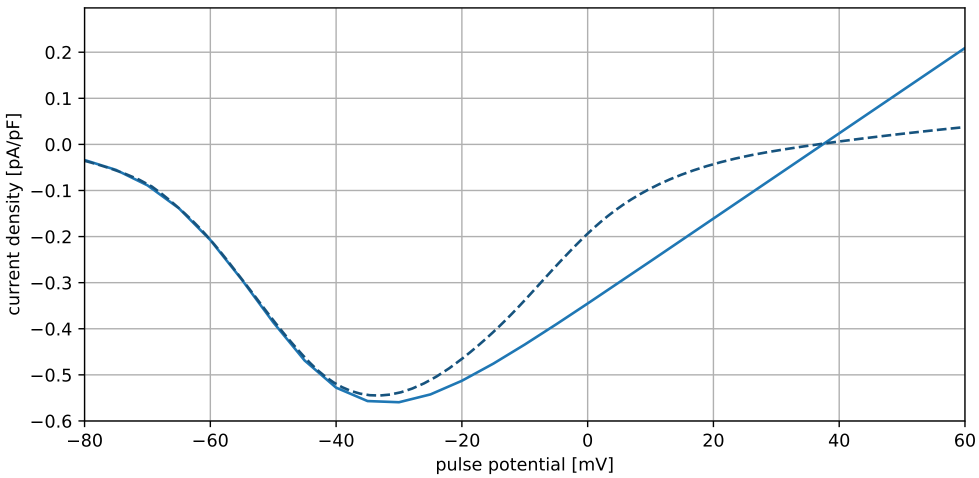

**Figure 20.** Current-voltage relationship of $I_{st}$ obtained with a voltage pulse protocol with a holding potential of -80 mV, a holding duration of 15 s, and a pulse duration of 500 ms. Solid line: Simulation result of SustainedInwardIV. Dashed line: Reference data extracted from Figure S5C in [1]. As in the previous figure, we had to use $g_{st} = 0.27$ nS to obtain a good agreement. Additionally, the behavior for positive pulse voltages is different. We do not have a good explanation for this difference, but can only point to the reference plot by Kurata2002 *et al.*, which shows a behavior that is more similar to InaMo than to the plot by Inada *et al.* (see Figure 30). This figure was created with InaMo version 1.4.2, which is available under the DOI 10.5281/zenodo.4533008.

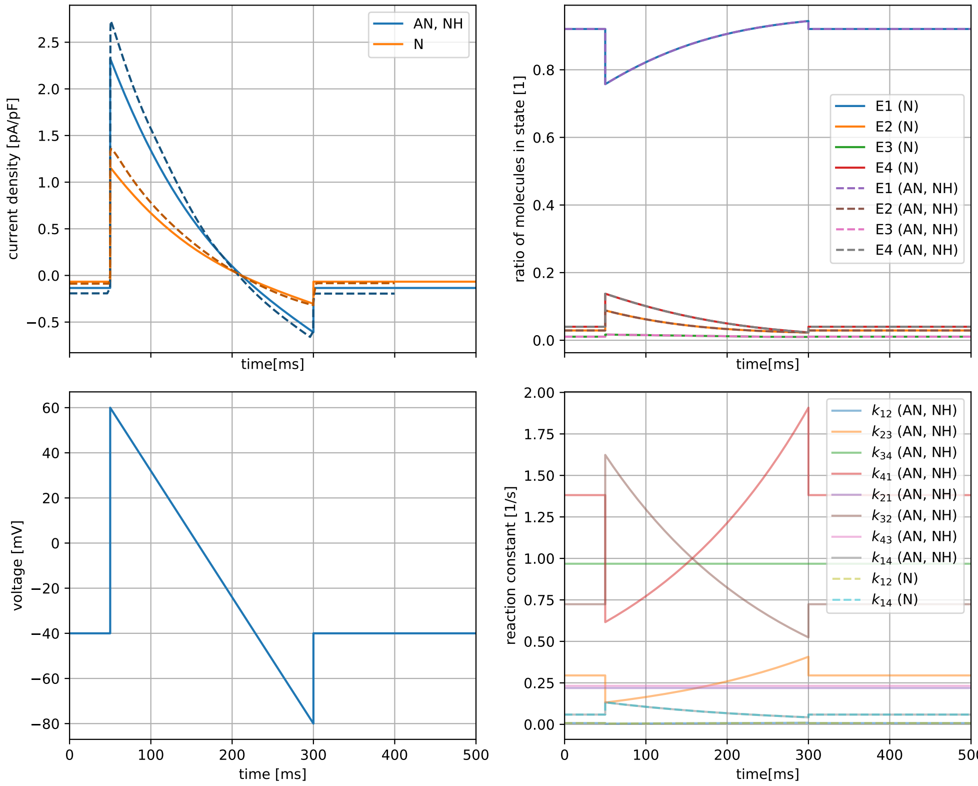

**Figure 21.** Current density time course of $I_{NaCa}$ following a ramp-shaped input voltage. Solid lines: Simulation result of SodiumCalciumExchangerRampInada. Dashed lines: Reference data extracted from Figure S6A in [1]. The current density in InaMo is slightly lower than in the reference. This can be explained by the fact that Inada *et al.* do not state whether calcium concentrations were held constant for the experiment and if so, which value was assumed for $\left[Ca^{2+}\right]_{sub}$. Since they used Data from Convery *et al.* [3], we assume that the calcium and sodium concentrations should be similar to those used in this experiment ($[Na^{+}]_i$ = 10 mM, $[Na^{+}]_o$ = 140 mM, $\left[Ca^{2+}\right]_o$ = 2.5 mM). However, Convery *et al.* do not give a value for $\left[Ca^{2+}\right]_{sub}$ and both using all values from Table S15 of [1] and using all values from [3] does not reproduce the absolute values observed in Figure S6. We therefore used a mix of settings from both sources and manually changed $\left[Ca^{2+}\right]_{sub}$ until a good fit with the reference was achieved. The remaining differences disappear when the current density is multiplied with a scaling factor of 1.18. This figure was created with InaMo version 1.4.2, which is available under the DOI 10.5281/zenodo.4533008.

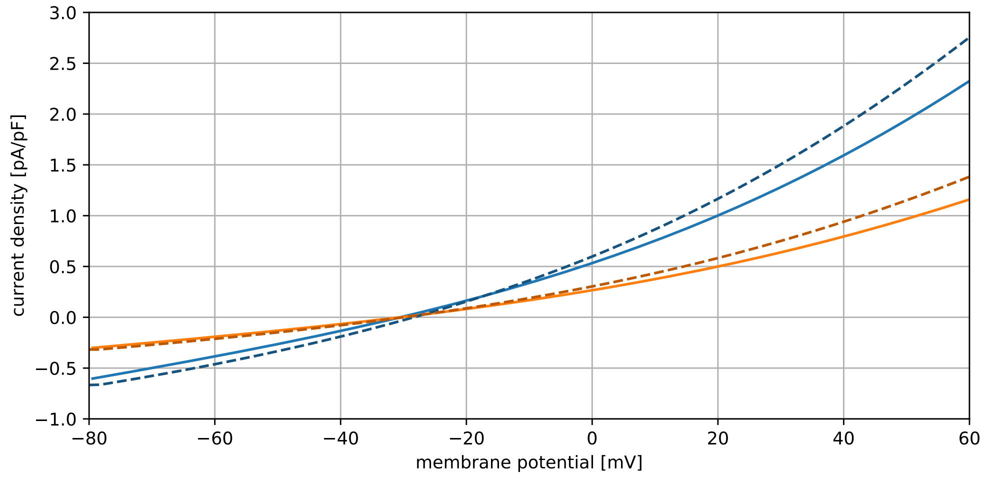

**Figure 22.** Current-voltage relationship of $I_{NaCa}$ obtained following a ramp-shaped input voltage (see previous figure). Solid lines: Simulation result of SodiumCalciumExchangerRampInada. Dashed lines: Reference data extracted from Figure S6B in [1]. As in the previous figure, InaMo has slightly lower current densities, which can be explained by missing information about the experiment setup of Inada *et al.*. Like in the previous Figure, the remaining differences disappear when the current density is multiplied with a scaling factor of 1.18. This figure was created with InaMo version 1.4.2, which is available under the DOI 10.5281/zenodo.4533008.

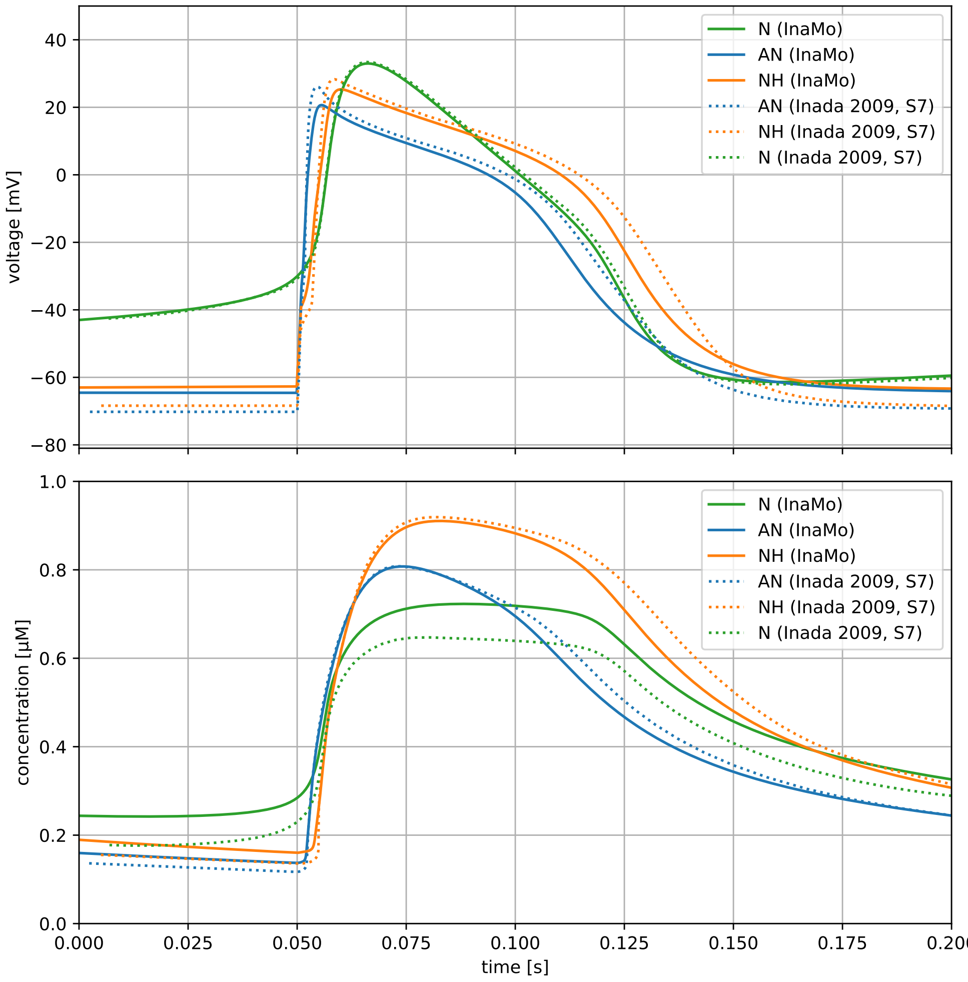

**Figure 23.** Time course of membrane voltage and $\left[Ca^{2+}\right]_i$ for the full cell models obtained by spontaneous activation (N cell) or stimulation with a current pulse protocol issuing pulses of -1.2 nA/-0.95 nA with a pulse duration of 1 ms after a holding period of 300 ms with a holding potential of 0 nA (AN/NH cell). Solid lines: Simulation result of AllCells. Dotted lines: Reference data extracted from Figure S7 in [1]. For AN and NH cells the voltage plot shows a lower resting potential and a slightly narrower action potential with respect to the reference. For N cells, $\left[Ca^{2+}\right]_i$ is significantly lower than in the reference. This can be explained by differences discussed in other figures. Additionally, the current pulse protocol is not given in [1]. We assume it is the same as for Figure 1 of [1], but this still only gives the stimulation current in terms of a multiplicative constant applied to an unspecified threshold current. We also changed the holding duration to 300 ms instead of 350 ms as it yields a better agreement. This figure was created with InaMo version 1.4.2, which is available under the DOI 10.5281/zenodo.4533008.

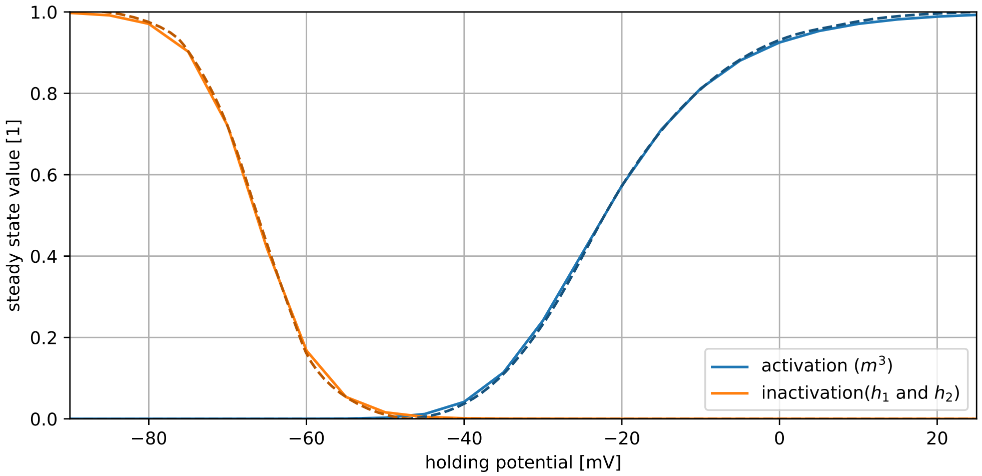

**Figure 24.** Steady state of gating variables for $I_{Na}$. Solid lines: Simulation result of SodiumChannelSteady. Dashed lines: Reference data extracted from Figure 2A in [4]. The plots are in perfect agreement. This figure was created with InaMo version 1.4.2, which is available under the DOI 10.5281/zenodo.4533008.

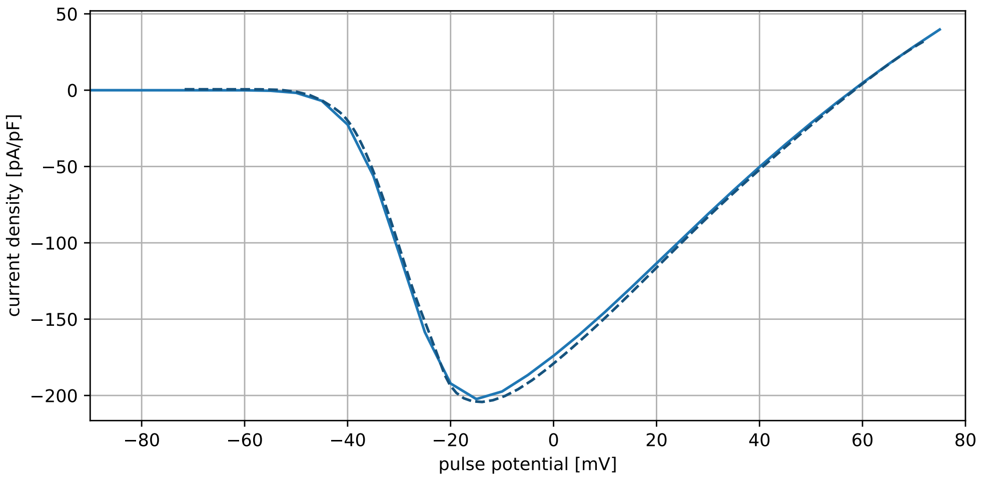

**Figure 25.** Current-voltage relationship of $I_{Na}$ obtained with a voltage pulse protocol with a holding potential of -90 mV, a holding duration of 2 s, and a pulse duration of 50 ms. Solid line: Simulation result of SodiumChannelIV. Dashed line: Reference data extracted from Figure 2B in [4]. The plots are in perfect agreement. This figure was created with InaMo version 1.4.2, which is available under the DOI 10.5281/zenodo.4533008.

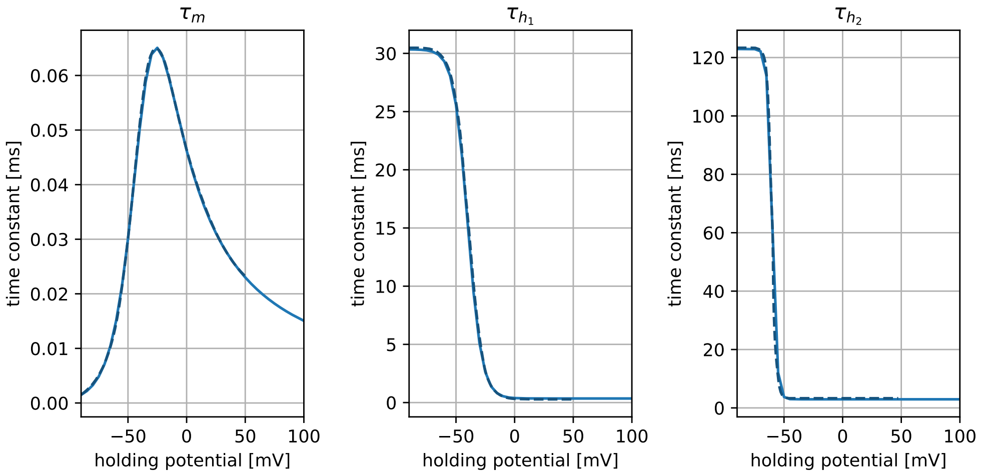

**Figure 26.** Time constants of gating variables for $I_{Na}$. Solid lines: Simulation result of SodiumChannelSteady. Dashed lines: Reference data extracted from Figures 2C–2E in [4]. The plots are in perfect agreement. This figure was created with InaMo version 1.4.2, which is available under the DOI 10.5281/zenodo.4533008.

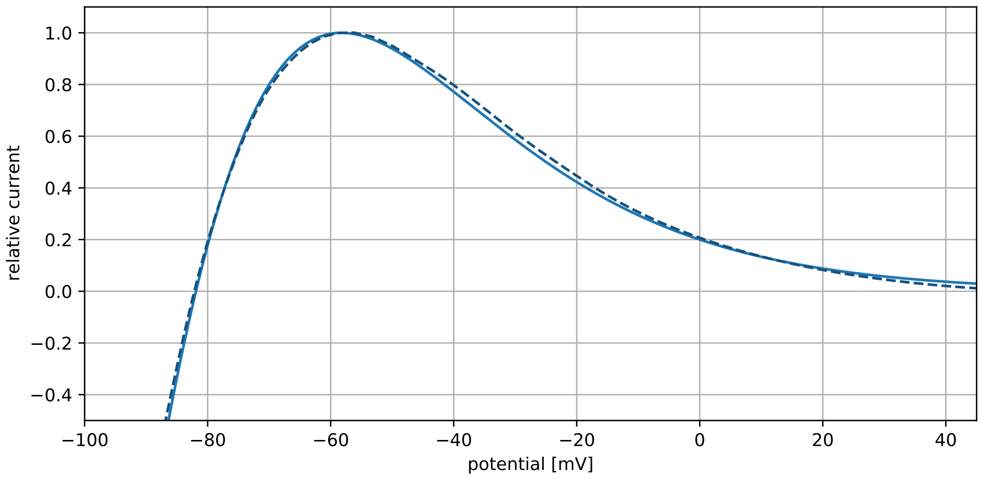

**Figure 27.** Current-voltage relationship of $I_{K,1}$ obtained by a voltage clamp experiment with linearly rising input voltage. Solid line: Simulation result of InwardRectifierLin. Dashed line: Reference data extracted from Figure 8 in [4]. The plots are in perfect agreement. This figure was created with InaMo version 1.4.2, which is available under the DOI 10.5281/zenodo.4533008.

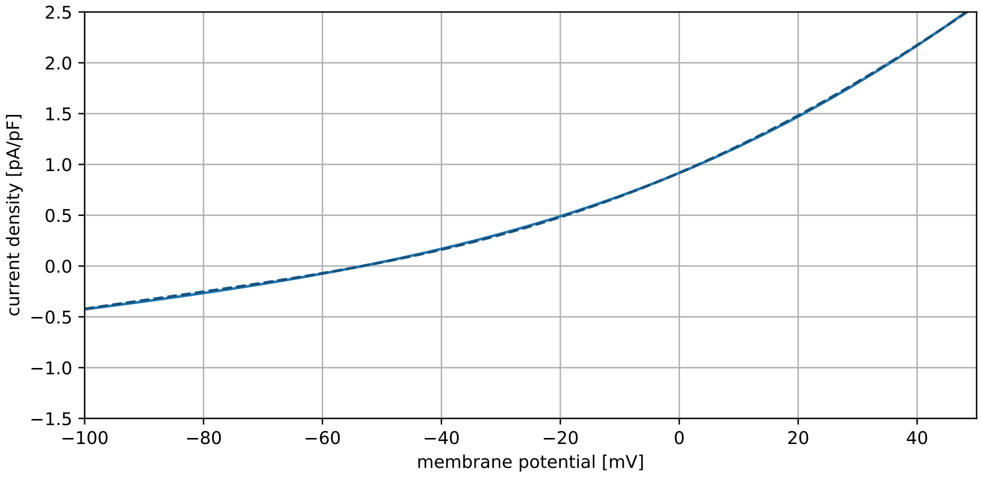

**Figure 28.** Current-voltage relationship of $I_{NaCa}$ obtained from a voltage clamp experiment with a linearly rising input voltage, which uses parameter settings from Kurata *et al.* [2]. Solid line: Simulation result of SodiumCalciumExchangerLinKurata. Dashed line: Reference data extracted from Figure 17 (upper left) in [2]. The plots are in perfect agreement. This figure was created with InaMo version 1.4.2, which is available under the DOI 10.5281/zenodo.4533008.

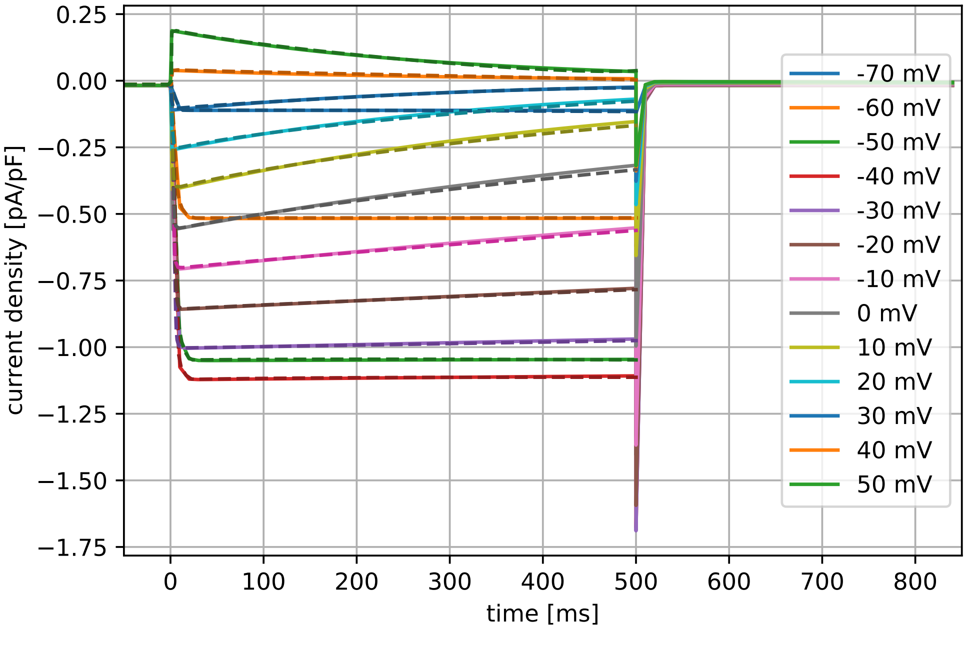

**Figure 29.** Current density time course of $I_{st}$ due to a 500 ms stimulation to different voltages after holding the voltage at -80 mV for 15 s. This experiment uses parameter settings and steady state equation from Kurata *et al.* [2]. Solid line: Simulation result of SustainedInwardIVKurata. Dashed line: Reference data extracted from Figure 4 (bottom left) in [2]. The plots are in perfect agreement. This figure was created with InaMo version 1.4.2, which is available under the DOI 10.5281/zenodo.4533008.

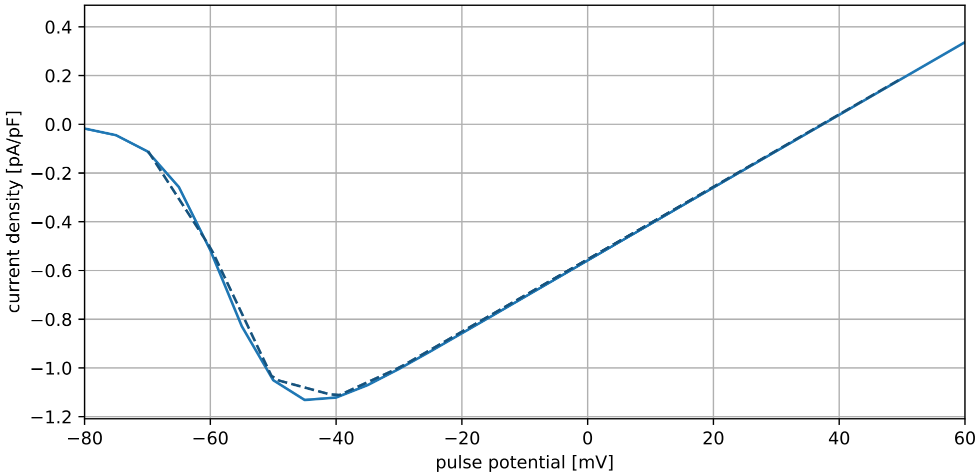

**Figure 30.** Current-voltage relationship of $I_{st}$ obtained with a voltage pulse protocol with a holding potential of -80 mV, a holding duration of 15 s, and a pulse duration of 500 ms. This experiment uses parameter settings and steady state equation from Kurata *et al.* [2]. Solid line: Simulation result of SustainedInwardIV. Dashed line: Reference data extracted from Figure 4 (bottom right) in [2]. The plots are in perfect agreement. This figure was created with InaMo version 1.4.2, which is available under the DOI 10.5281/zenodo.4533008.

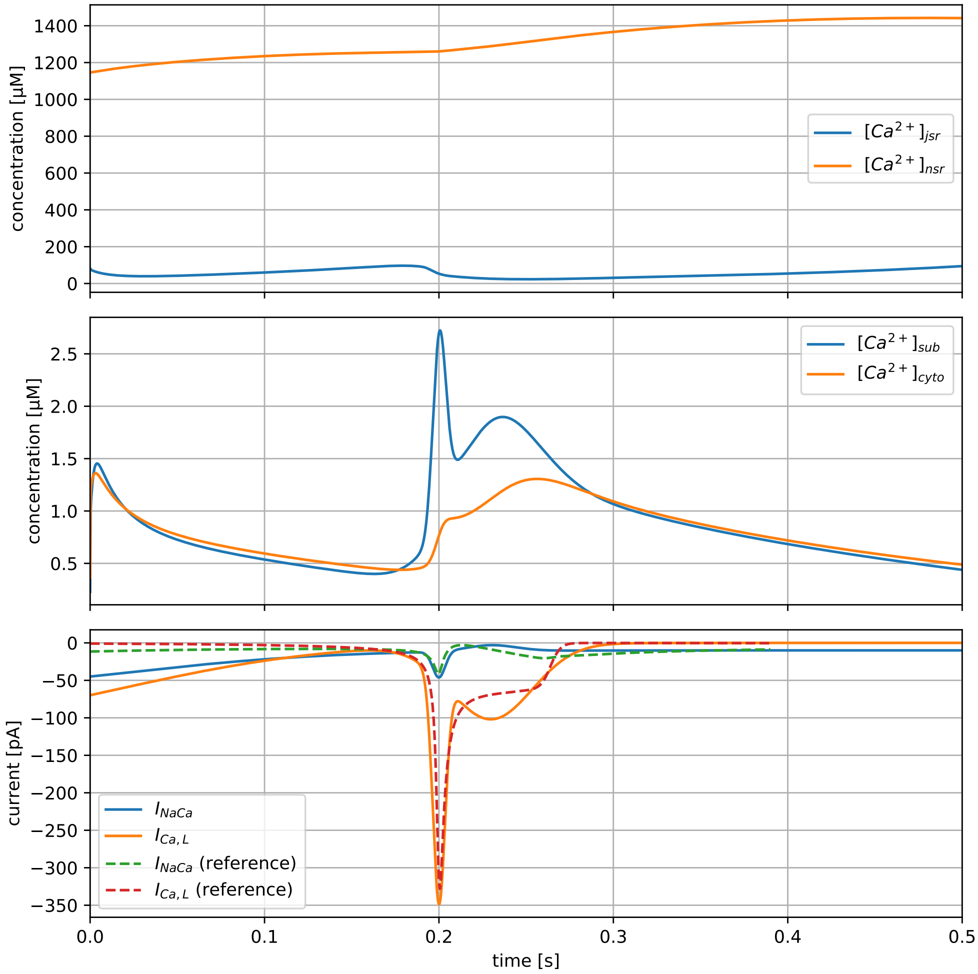

**Figure 31.** Time course of $Ca^{2+}$ concentrations due to a dummy current for $I_{NaCa}$ and $I_{Ca,L}$, which roughly resemble the true currents during an action potential (dotted lines in bottom plot). No reference plot is available for this experiment, but it allows to examine the $\left[Ca^{2+}\right]$ handling in isolation from other components. This figure was created with InaMo version 1.4.2, which is available under the DOI 10.5281/zenodo.4533008.

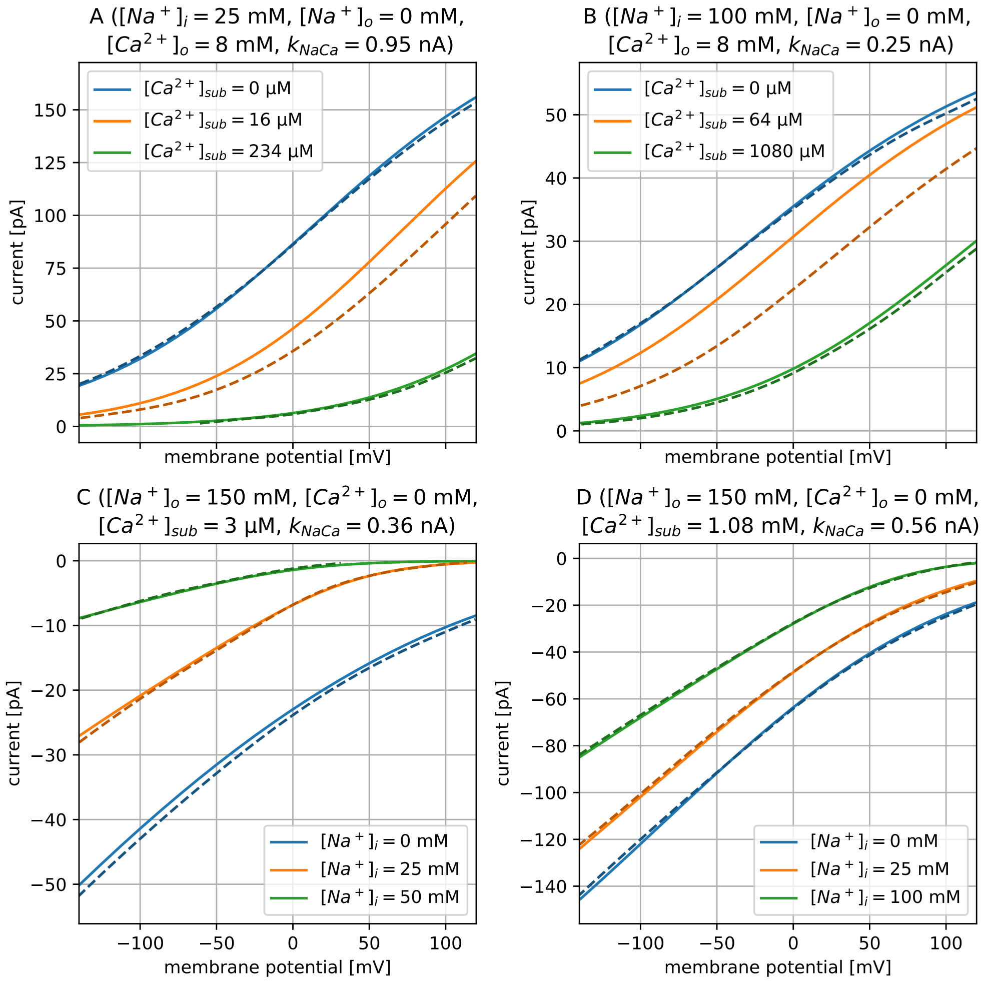

**Figure 32.** Current-voltage relationship of $I_{NaCa}$ obtained from a voltage clamp experiment with linearly rising input voltage. Parts A and B show effect of variation in $[Ca^{2+}]_{sub}$ and parts C and D show effect of variation in $[Na^{+}]_{i}$. Solid lines: Simulation result of SodiumCalciumExchangerLinMatsuoka. Dashed lines: Reference data extracted from Figure 19 in [5]. Absolute current values are not shown in Figure 19, but can be found in Figures 15A (A), Figure 16B (B), and Figure 17 (C, D) in [5]. For part A and B, the lines with 0 and 16  M (A) and 0 and 64  M (B) are closer to each other, which does not agree with the reference, but does agree with the experimental data (filled and open circles). We have no good explanation for this difference, since it only occurs in two of the four experiment setups. The absolute values are not exact, since Matsuoka *et al.* used a different scaling factor for each part but do not report its value. We therefore choose the value of $k_{NaCa}$ freely to roughly reproduce the absolute values in Figures 15–17 in [5]. This figure was created with InaMo version 1.4.2, which is available under the DOI 10.5281/zenodo.4533008.

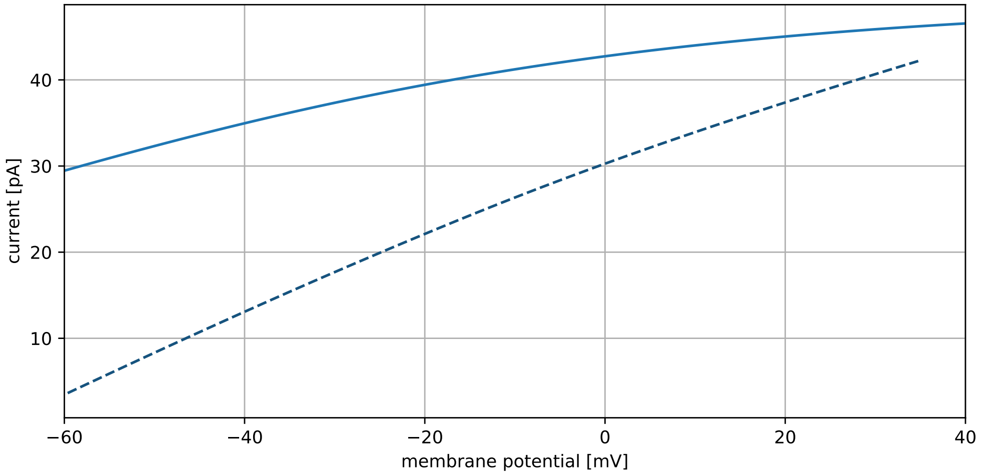

**Figure 33.** Current-voltage relationship of $I_p$ obtained from a voltage clamp experiment with linearly rising input voltage. Solid line: Simulation result of SodiumPotassiumPumpLin. Dashed line: Reference data extracted from Figure 12 in [6]. The reference cannot be reproduced exactly as Demir *et al.* only report the sum of $I_p$ and three different background currents. This figure was created with InaMo version 1.4.2, which is available under the DOI 10.5281/zenodo.4533008.

### 2 Supplementary Tables

| Model Part | Variable | Unit | Article | C++ | CellML |
|---|---|---|---|---|---|
| Membrane | $V$ | V | $-7.003 \cdot 10^{-2}$ | $-7.022 \cdot 10^{-2}$ | $-7.155 \cdot 10^{-2}$ |
| $I_{Na}$ | $m$ | 1 | $1.227 \cdot 10^{-2}$ | $1.246 \cdot 10^{-2}$ | $1.048 \cdot 10^{-2}$ |
| $I_{Na}$ | $h_1$ | 1 | $7.170 \cdot 10^{-1}$ | $7.273 \cdot 10^{-1}$ | $7.922 \cdot 10^{-1}$ |
| $I_{Na}$ | $h_2$ | 1 | $6.162 \cdot 10^{-1}$ | $6.378 \cdot 10^{-1}$ | $7.883 \cdot 10^{-1}$ |
| $I_{Ca,L}$ | $d_L$ | 1 | $4.069 \cdot 10^{-5}$ | $3.952 \cdot 10^{-5}$ | $3.229 \cdot 10^{-5}$ |
| $I_{Ca,L}$ | $f_{L,\text{fast}}$ | 1 | $9.985 \cdot 10^{-1}$ | $9.985 \cdot 10^{-1}$ | $9.988 \cdot 10^{-1}$ |
| $I_{Ca,L}$ | $f_{L,\text{slow}}$ | 1 | $9.875 \cdot 10^{-1}$ | $9.911 \cdot 10^{-1}$ | $9.988 \cdot 10^{-1}$ |
| $I_{to}$ | $r$ | 1 | $8.857 \cdot 10^{-3}$ | $8.704 \cdot 10^{-3}$ | $8.029 \cdot 10^{-3}$ |
| $I_{to}$ | $q_{\text{fast}}$ | 1 | $8.734 \cdot 10^{-1}$ | $8.847 \cdot 10^{-1}$ | $9.955 \cdot 10^{-1}$ |
| $I_{to}$ | $q_{\text{slow}}$ | 1 | $1.503 \cdot 10^{-1}$ | $2.026 \cdot 10^{-1}$ | $5.480 \cdot 10^{-1}$ |
| $I_{K,r}$ | $p_{\text{a, fast}}$ | 1 | $7.107 \cdot 10^{-2}$ | $3.473 \cdot 10^{-2}$ | $9.078 \cdot 10^{-4}$ |
| $I_{K,r}$ | $p_{\text{a, slow}}$ | 1 | $4.840 \cdot 10^{-2}$ | $3.473 \cdot 10^{-2}$ | $2.899 \cdot 10^{-3}$ |
| $I_{K,r}$ | $p_{\text{i}}$ | 1 | $9.866 \cdot 10^{-1}$ | $9.868 \cdot 10^{-1}$ | $9.879 \cdot 10^{-1}$ |
| $[Ca^{2+}]$ handling | $[Ca^{2+}]_i$ | mM | $1.206 \cdot 10^{-4}$ | $1.082 \cdot 10^{-4}$ | $3.104 \cdot 10^{-5}$ |
| $[Ca^{2+}]$ handling | $[Ca^{2+}]_{sub}$ | mM | $6.397 \cdot 10^{-5}$ | $5.860 \cdot 10^{-5}$ | $2.870 \cdot 10^{-5}$ |
| $[Ca^{2+}]$ handling | $[Ca^{2+}]_{jsr}$ | mM | $4.273 \cdot 10^{-1}$ | $4.002 \cdot 10^{-1}$ | $5.575 \cdot 10^{-1}$ |
| $[Ca^{2+}]$ handling | $[Ca^{2+}]_{nsr}$ | mM | $1.068 \cdot 10^{+0}$ | $9.596 \cdot 10^{-1}$ | $6.672 \cdot 10^{-1}$ |
| $[Ca^{2+}]$ handling | $f_{TC}$ | 1 | $2.359 \cdot 10^{-2}$ | $2.120 \cdot 10^{-2}$ | $6.156 \cdot 10^{-3}$ |
| $[Ca^{2+}]$ handling | $f_{TMC}$ | 1 | $3.667 \cdot 10^{-1}$ | $3.395 \cdot 10^{-1}$ | $1.122 \cdot 10^{-1}$ |
| $[Ca^{2+}]$ handling | $f_{TMM}$ | 1 | $5.594 \cdot 10^{-1}$ | $5.834 \cdot 10^{-1}$ | $7.843 \cdot 10^{-1}$ |
| $[Ca^{2+}]$ handling | $f_{CM_i}$ | 1 | $4.845 \cdot 10^{-2}$ | $4.366 \cdot 10^{-2}$ | $1.290 \cdot 10^{-2}$ |
| $[Ca^{2+}]$ handling | $f_{CM_s}$ | 1 | $2.626 \cdot 10^{-2}$ | $2.410 \cdot 10^{-2}$ | $1.192 \cdot 10^{-2}$ |
| $[Ca^{2+}]$ handling | $f_{CQ}$ | 1 | $3.379 \cdot 10^{-1}$ | $3.235 \cdot 10^{-1}$ | $4.007 \cdot 10^{-1}$ |
| $[Ca^{2+}]$ handling | $f_{CSL}$ | 1 | $3.936 \cdot 10^{-5}$ | $3.085 \cdot 10^{-5}$ | $8.911 \cdot 10^{-6}$ |

**Table 1.** Initial values for AN cell model in article, C++-code, and CellML code.

| Model Part | Variable | Unit | Article | C++ | CellML |
|---|---|---|---|---|---|
| Membrane | $V$ | V | $-6.213 \cdot 10^{-2}$ | $-6.213 \cdot 10^{-2}$ | $-4.971 \cdot 10^{-2}$ |
| $I_{Ca,L}$ | $d_L$ | 1 | $1.533 \cdot 10^{-4}$ | $1.534 \cdot 10^{-4}$ | $1.793 \cdot 10^{-3}$ |
| $I_{Ca,L}$ | $f_{L,\text{fast}}$ | 1 | $6.861 \cdot 10^{-1}$ | $6.809 \cdot 10^{-1}$ | $9.756 \cdot 10^{-1}$ |
| $I_{Ca,L}$ | $f_{L,\text{slow}}$ | 1 | $4.441 \cdot 10^{-1}$ | $3.320 \cdot 10^{-1}$ | $7.744 \cdot 10^{-1}$ |
| $I_{K,r}$ | $p_{\text{a, fast}}$ | 1 | $6.067 \cdot 10^{-1}$ | $6.061 \cdot 10^{-1}$ | $1.925 \cdot 10^{-1}$ |
| $I_{K,r}$ | $p_{\text{a, slow}}$ | 1 | $1.287 \cdot 10^{-1}$ | $1.288 \cdot 10^{-1}$ | $7.972 \cdot 10^{-2}$ |
| $I_{K,r}$ | $p_{\text{i}}$ | 1 | $9.775 \cdot 10^{-1}$ | $9.775 \cdot 10^{-1}$ | $9.490 \cdot 10^{-1}$ |
| $I_f$ | $y$ | 1 | $3.825 \cdot 10^{-2}$ | $3.823 \cdot 10^{-2}$ | $4.623 \cdot 10^{-2}$ |
| $I_{st}$ | $q_a$ | 1 | $1.933 \cdot 10^{-1}$ | $1.936 \cdot 10^{-1}$ | $4.764 \cdot 10^{-1}$ |
| $I_{st}$ | $q_i$ | 1 | $4.886 \cdot 10^{-1}$ | $4.885 \cdot 10^{-1}$ | $5.423 \cdot 10^{-1}$ |
| $[Ca^{2+}]$ handling | $[Ca^{2+}]_i$ | mM | $3.623 \cdot 10^{-4}$ | $3.633 \cdot 10^{-4}$ | $1.850 \cdot 10^{-4}$ |
| $[Ca^{2+}]$ handling | $[Ca^{2+}]_{sub}$ | mM | $2.294 \cdot 10^{-4}$ | $2.300 \cdot 10^{-4}$ | $1.603 \cdot 10^{-4}$ |
| $[Ca^{2+}]$ handling | $[Ca^{2+}]_{jsr}$ | mM | $8.227 \cdot 10^{-2}$ | $8.185 \cdot 10^{-2}$ | $2.963 \cdot 10^{-1}$ |
| $[Ca^{2+}]$ handling | $[Ca^{2+}]_{nsr}$ | mM | $1.146 \cdot 10^{+0}$ | $1.146 \cdot 10^{+0}$ | $1.111 \cdot 10^{+0}$ |
| $[Ca^{2+}]$ handling | $f_{TC}$ | 1 | $6.838 \cdot 10^{-1}$ | $6.856 \cdot 10^{-2}$ | $3.565 \cdot 10^{-2}$ |
| $[Ca^{2+}]$ handling | $f_{TMC}$ | 1 | $6.192 \cdot 10^{-1}$ | $6.195 \cdot 10^{-1}$ | $4.433 \cdot 10^{-1}$ |
| $[Ca^{2+}]$ handling | $f_{TMM}$ | 1 | $3.363 \cdot 10^{-1}$ | $3.360 \cdot 10^{-1}$ | $3.917 \cdot 10^{-1}$ |
| $[Ca^{2+}]$ handling | $f_{CM_i}$ | 1 | $1.336 \cdot 10^{-1}$ | $1.339 \cdot 10^{-1}$ | $7.230 \cdot 10^{-2}$ |
| $[Ca^{2+}]$ handling | $f_{CM_s}$ | 1 | $8.894 \cdot 10^{-2}$ | $8.915 \cdot 10^{-2}$ | $6.308 \cdot 10^{-2}$ |
| $[Ca^{2+}]$ handling | $f_{CQ}$ | 1 | $8.736 \cdot 10^{-2}$ | $8.694 \cdot 10^{-2}$ | $2.614 \cdot 10^{-1}$ |
| $[Ca^{2+}]$ handling | $f_{CSL}$ | 1 | $4.764 \cdot 10^{-5}$ | $4.674 \cdot 10^{-5}$ | $4.150 \cdot 10^{-5}$ |

**Table 2.** Initial values for N cell model in article, C++ code, and CellML code.

| Model Part | Variable | Unit | Article | C++ | CellML |
|---|---|---|---|---|---|
| Membrane | $V$ | V | $-6.863 \cdot 10^{-2}$ | $-6.867 \cdot 10^{-2}$ | $-6.976 \cdot 10^{-2}$ |
| $I_{Na}$ | $m$ | 1 | $1.529 \cdot 10^{-2}$ | $1.521 \cdot 10^{-2}$ | $1.322 \cdot 10^{-2}$ |
| $I_{Na}$ | $h_1$ | 1 | $6.438 \cdot 10^{-1}$ | $6.463 \cdot 10^{-1}$ | $7.066 \cdot 10^{-1}$ |
| $I_{Na}$ | $h_2$ | 1 | $5.552 \cdot 10^{-1}$ | $5.638 \cdot 10^{-1}$ | $7.016 \cdot 10^{-1}$ |
| $I_{Ca,L}$ | $d_L$ | 1 | $5.025 \cdot 10^{-5}$ | $4.996 \cdot 10^{-5}$ | $4.235 \cdot 10^{-5}$ |
| $I_{Ca,L}$ | $f_{L,\text{fast}}$ | 1 | $9.981 \cdot 10^{-1}$ | $9.981 \cdot 10^{-1}$ | $9.984 \cdot 10^{-1}$ |
| $I_{Ca,L}$ | $f_{L,\text{slow}}$ | 1 | $9.831 \cdot 10^{-1}$ | $9.851 \cdot 10^{-1}$ | $9.984 \cdot 10^{-1}$ |
| $I_{to}$ | $r$ | 1 | $9.581 \cdot 10^{-3}$ | $9.559 \cdot 10^{-3}$ | $8.948 \cdot 10^{-3}$ |
| $I_{to}$ | $q_{\text{fast}}$ | 1 | $8.640 \cdot 10^{-1}$ | $8.708 \cdot 10^{-1}$ | $9.948 \cdot 10^{-1}$ |
| $I_{to}$ | $q_{\text{slow}}$ | 1 | $1.297 \cdot 10^{-1}$ | $1.343 \cdot 10^{-1}$ | $4.274 \cdot 10^{-1}$ |
| $I_{K,r}$ | $p_{\text{a, fast}}$ | 1 | $9.949 \cdot 10^{-2}$ | $9.333 \cdot 10^{-2}$ | $1.418 \cdot 10^{-3}$ |
| $I_{K,r}$ | $p_{\text{a, slow}}$ | 1 | $7.024 \cdot 10^{-2}$ | $6.769 \cdot 10^{-2}$ | $5.398 \cdot 10^{-3}$ |
| $I_{K,r}$ | $p_{\text{i}}$ | 1 | $9.853 \cdot 10^{-1}$ | $9.854 \cdot 10^{-1}$ | $9.864 \cdot 10^{-1}$ |
| $\left[Ca^{2+}\right]$ handling | $\left[Ca^{2+}\right]_i$ | mM | $1.386 \cdot 10^{-4}$ | $1.340 \cdot 10^{-4}$ | $3.733 \cdot 10^{-5}$ |
| $\left[Ca^{2+}\right]$ handling | $\left[Ca^{2+}\right]_{sub}$ | mM | $7.314 \cdot 10^{-5}$ | $7.122 \cdot 10^{-5}$ | $3.273 \cdot 10^{-5}$ |
| $\left[Ca^{2+}\right]$ handling | $\left[Ca^{2+}\right]_{jsr}$ | mM | $4.438 \cdot 10^{-1}$ | $4.475 \cdot 10^{-1}$ | $6.822 \cdot 10^{-1}$ |
| $\left[Ca^{2+}\right]$ handling | $\left[Ca^{2+}\right]_{nsr}$ | mM | $1.187 \cdot 10^{+0}$ | $1.163 \cdot 10^{+0}$ | $8.187 \cdot 10^{-1}$ |
| $\left[Ca^{2+}\right]$ handling | $f_{TC}$ | 1 | $2.703 \cdot 10^{-2}$ | $2.615 \cdot 10^{-2}$ | $7.396 \cdot 10^{-3}$ |
| $\left[Ca^{2+}\right]$ handling | $f_{TMC}$ | 1 | $4.020 \cdot 10^{-1}$ | $3.930 \cdot 10^{-1}$ | $1.337 \cdot 10^{-1}$ |
| $\left[Ca^{2+}\right]$ handling | $f_{TMM}$ | 1 | $5.282 \cdot 10^{-1}$ | $5.362 \cdot 10^{-1}$ | $7.652 \cdot 10^{-1}$ |
| $\left[Ca^{2+}\right]$ handling | $f_{CM_i}$ | 1 | $5.530 \cdot 10^{-2}$ | $5.355 \cdot 10^{-2}$ | $1.547 \cdot 10^{-2}$ |
| $\left[Ca^{2+}\right]$ handling | $f_{CM_s}$ | 1 | $2.992 \cdot 10^{-2}$ | $2.911 \cdot 10^{-2}$ | $1.358 \cdot 10^{-2}$ |
| $\left[Ca^{2+}\right]$ handling | $f_{CQ}$ | 1 | $3.463 \cdot 10^{-1}$ | $3.483 \cdot 10^{-1}$ | $4.500 \cdot 10^{-1}$ |
| $\left[Ca^{2+}\right]$ handling | $f_{CSL}$ | 1 | $4.843 \cdot 10^{-5}$ | $4.447 \cdot 10^{-5}$ | $1.217 \cdot 10^{-5}$ |

**Table 3.** Initial values for NH cell model in article, C++-code, and CellML code.
